# Structural determinants of rotavirus proteolytic activation

**DOI:** 10.1101/2025.03.24.644915

**Authors:** Dunia Asensio-Cob, Carlos P. Mata, Josué Gómez-Blanco, Javier Vargas, Javier M. Rodríguez, Daniel Luque

**Affiliations:** Department of Molecular Medicine, Peter Gilgan Centre for Research and Learning, The Hospital for Sick Children, 686 Bay Street, Toronto ON, M5G0A4, Canada; Departamento de Estructura de Macromoléculas, Centro Nacional de Biotecnología, CSIC, Cantoblanco, 28049 Madrid, Spain; Artis Analytica Scientia S.L., Madrid 28021, Spain; Departamento de Óptica, Universidad Complutense de Madrid, Madrid 28040, Spain; Electron Microscope Unit, Mark Wainwright Analytical Centre, University of New South Wales, Sydney, NSW 2052, Australia; School of Biomedical Sciences, University of New South Wales, Sydney, NSW 2052, Australia

**Author notes:** These authors contributed equally to this work (Dunia Asensio-Cob, Carlos P. Mata). Co-corresponding authors: Javier M. Rodríguez, and Daniel Luque.

**Keywords:** Rotavirus, viral entry, *Reovirales*, cryo-electron microscopy, proteolytic activation

## Abstract

The infectivity of rotavirus (RV), the leading cause of childhood diarrhea, hinges on the activation of viral particles through the proteolysis of the spike protein by trypsin-like proteases in the host intestinal lumen. Despite comprehensive structural characterization of the virus particle, the structural rationale behind the necessity of trypsin digestion of the VP4 protein for infectivity remains poorly understood. In this study, using cryo-electron microscopy (cryo-EM) and advanced image processing techniques, we compared uncleaved and cleaved RV virions and found that the conformation of the non-proteolyzed spike is constrained by the position of loops that surround its structure, linking the lectin domains of the spike head to its body. The proteolysis of these loops removes this structural constraint, thereby enabling the spike to undergo the necessary conformational changes required for cell membrane penetration. Thus, these loops function as regulatory elements to ensure that the spike protein is activated precisely when and where it is needed to facilitate a successful infection.

## Introduction

During the viral replication cycle, entry emerges as the first critical juncture facing the viral particle for successful infection. The processes involved are tightly regulated in both space and time, with success only occurring after the carefully orchestrated initial stages of infection have been successfully completed. For some viruses, these early infection steps lead to an irreversible commitment to entry, which, if unsuccessful, renders the particle non-infectious. Consequently, these viruses have evolved sensing mechanisms to ensure that entry is only attempted in the correct environment. Many viral proteins responsible for crossing the host cell membrane are synthesized as inactive precursors that undergo proteolytic cleavage to become fully active (Forzan et al., 2004; Izaguirre, 2019; Ykema et al., 2023). Processing of viral surface proteins by host proteases, like furin and trypsin, is a control mechanism that has been shown to produce mature virions activated for infection in a wide range of viruses. While, typically, proteolytic activation allows membrane fusion of enveloped virus surface proteins (Bertram et al., 2011; Bestle et al., 2021; Krzyzaniak et al., 2013; Menachery et al., 2020; Tang et al., 2021; White & Whittaker, 2016), in many non-enveloped viruses, is essential for membrane penetration (Aguilera-Flores et al., 2022; Crawford et al., 2001; Graham & Estes, 1980; Johnson & Banerjee, 2008; Pletan & Tsai, 2022; Ykema et al., 2023). Astroviruses and rotaviruses (RV) are paradigmatic examples of non-enveloped enteric viruses whose infectivity is dependent on trypsin-like digestive enzymes present in the gastrointestinal tract (Bass & Qiu, 2000; Clark et al., 1981; Graham & Estes, 1980; Lee & Kurtz, 1981; Ludert et al., 1996).

RV are non-enveloped, double-stranded RNA (dsRNA) viruses of the order *Reovirales* that replicate in differentiated enterocytes of the intestinal tract in vertebrates, and are transmitted predominantly via the faecal-oral route (Howley et al., 2023). The RV virion is an icosahedral triple layered particle in which the segmented viral genome and the polymerase complexes are packaged in a capsid made of the VP2 protein (McClain et al., 2010; Settembre et al., 2011) (<u>Fig. S1A</u>). This core is surrounded by an intermediate layer of the VP6 protein trimers, originating the double-layered particle (DLP) (Mathieu et al., 2001; McClain et al., 2010). The external layer is formed by trimers of the VP7 glycoprotein in phase with the underlaying VP6 shell and 60 trimeric VP4 spikes, which project from the VP7 layer, that clamps them and partially covers their base (Herrmann et al., 2021; Shah et al., 2023). This outer layer is responsible for the first steps of RV infection: attachment to cell surface receptors, lipid bilayer penetration, and release of the double-layered particle (DLP) into the cytosol that would start the genetic program of the virus (Abdelhakim et al., 2014; C. F. Arias & López, 2021; Herrmann et al., 2021).

During RV morphogenesis, the nascent DLP exits the viroplasm just before budding into the ER lumen through interactions of the VP6 layer with the viral NSP4 protein, located in the ER membrane (Au et al., 1993; Taylor et al., 1993), resulting in a transiently enveloped DLP (eDLP). VP4 interacts with both NSP4 and VP6 in the interphase between the viroplasm and the ER, and is incorporated into the eDLP (Au et al., 1993; Shah et al., 2023). In the eDLP, VP4 spikes are assembled as pre-mature symmetric trimers anchored onto the VP6 hexameric cavities surrounding the five-fold axes through a ‘foot’ contributed by the C-terminal regions of the three chains (<u>Fig. S1B-C</u>). Three spaced lobes, constituted by the VP4 β-barrel and lectin domains, protrude in the space between the DLP and the membrane of the eDLP (Shah et al., 2023). It has been proposed that a rapid process, which includes the extension of VP4 to breach the membrane followed by the assembly of the VP7 layer, will lead to the loss of the transient envelope and the formation of immature TLP in the ER lumen (Shah et al., 2023).

In the spike of the non-infectious immature TLP, the three VP4 molecules, VP4A, –B, and –C, are organized into a complex asymmetric conformation held by non-covalent interactions among its chains (Herrmann et al., 2021; Shah et al., 2023). While the C-terminal region of the three chains maintain a three-fold symmetry association into the ‘foot’ of the spike, the extension of the pre-mature VP4 lobes generates an upright asymmetric ‘dimeric’ conformation (<u>Fig. S1C</u>). The ‘stalk’ region contains the β-barrel domain from the C subunit, while the equivalent β-barrels from the A and B subunits form the dimeric ‘body’ of the spike. The lectin domains from VP4A and –B form the spike ‘head,’ which covers the hydrophobic loops of the two body β-barrels, while the third VP4-C lectin domain is prone and located at the spike stalk. Newly assembled immature TLP exit the cell either through exocytosis or lysis, depending on the specific cell line (Iša et al., 2020; Jourdan et al., 1997; Kerviel et al., 2021; Musalem & Espejo, 1985). Once in the intestinal lumen, the proteolytic processing of the virions by trypsin-like proteases result in infectious mature TLP (Clark et al., 1981; Crawford et al., 2001; Graham & Estes, 1980). In vitro, trypsin allows rotavirus infectivity and led to the activation of the RV spike by cleaving the VP4 protein after three defined sites (Arg231, Arg241, and Arg247) into an N-terminal fragment, VP8*, and a C-terminal fragment, VP5* (<u>Fig. S1C</u>). After trypsin cleavage, VP8*-A and –B lectin domains remain associated within the spike head but the stalk VP8*-C lectin domain is lost (Settembre et al., 2011). It has been suggested that an additional trypsin cleavage at Lys29, facilitates the removal of VP8*-C lectin domain (Settembre et al., 2011).

After activation by trypsin digestion, mature TLP initiates infection by binding of VP8* to its cellular receptor, which promotes TLP internalization either by virus-directed engulfment (Abdelhakim et al., 2014) or endocytosis (C. F. Arias et al., 2015; Diaz-Salinas et al., 2014). The attachment of VP8* initiates a sequence of structural changes that will ultimately facilitate membrane penetration and the delivery of DLP into the host cell cytoplasm (de Sautu et al., 2024; PR. Dormitzer et al., 2004; Herrmann et al., 2021; Trask et al., 2010; Wolf et al., 2012; Yoder et al., 2009). In the initial step, VP8* head lectin domains move laterally, allowing the spike to transition from an asymmetric ‘upright’ conformation to a symmetric ‘intermediate’ conformation (<u>Fig. S1C</u>). In this intermediate conformation, the hydrophobic loops of the three VP5* β-barrel interact with the lipid bilayer of the target membrane (PR. Dormitzer et al., 2004). The interaction between the hydrophobic loops and the membrane then promotes a ‘reversed’ conformation involving the unfolding and outward extension of the three VP5* foot domains, which perforate and penetrate the lipid bilayer of the vesicle membrane (<u>Fig. S1C</u>).

Despite the detailed structural information that has allowed the identification of the different conformational stages that the spike protein transits from its assembly into eDLP in the ER to its dissociation during membrane penetration, the structural basis underlaying the proteolytic activation from an immature to a mature spike remains unclear. Here, we use cryo-EM and computational methods to compare the structure of the uncleaved and cleaved spike of rotavirus. The resolved structures show that in the non-trypsinized spike the lateral movement of the lectin domains is restricted by the position of the loops that surround its structure and connect the lectin domains of the head with its body. Proteolysis of these loops breaks this structural constraint, allowing the necessary conformational changes to penetrate the cell membrane.

## Results

Previous structural cryo-EM characterizations of immature (Rodríguez et al., 2014; Shah et al., 2023) and mature TLP (Settembre et al., 2011) have revealed that the larger difference between uncleaved and cleaved spikes lies in the presence or absence of the VP4-C head domain at the base of the spike. However, it is unclear how the release of this domain could enhance the particle infectivity. On the other hand, the structure of the loop connecting the head and body domains of VP4 remains elusive. This region contains three trypsin targets (Arg231, Arg241, and Arg247) that are processed in a defined order (C. Arias et al., 1996) to generate an infectious RV particle. While the icosahedral symmetry of the RV capsid has facilitated the structural study of the VP2, VP6 and VP7 layers using cryo-EM and single particle analysis (SPA), it complicates the study of the spike at high-resolution due to its flexible nature and the partial depletion of spikes around the RV particle (D. Chen & Ramig, 1992; Ilca et al., 2015; Rodríguez et al., 2014; Settembre et al., 2011; Trask & Dormitzer, 2006). By combining SPA with advanced image processing methods (as described by (Ilca et al., 2015; Kaur et al., 2021; Kazemi et al., 2021), we aimed to address the continuous flexibility and partial ocupancy of the spike.

## Single particle analysis of NTR– and TR-TLP

To obtain high-resolution maps of activated and inactivated particles, TLP from cells infected with rotavirus in the presence of trypsin (TR-TLP), or in its absence in medium supplemented with the protease inhibitor leupeptin (NTR-TLP), were purified by density gradient ultracentrifugation and characterized by SDS-PAGE (<u>Fig. S2</u>). A single band corresponding to the 98 kDa unprocessed precursor form, VP4, was detected for the NTR-TLP spike protein. In contrast, for the TR-TLP, the proteolytic products VP8* (28 kDa) and VP5* (55 kDa) were detected as two distinct bands. Analysis of the particles by cryo-EM shows a homogeneous population of isometric TLP, some of which can be seen with VP4 (or VP8*/VP5*) spikes projected from the TLP surface (<u>Fig. S2, arrowheads</u>). Three-dimensional reconstructions (3DR) of NTR– (Fig. 1A-C) and TR-TLP (Fig. 1D-F) were computed using SPA with imposition of icosahedral symmetry, achieving resolutions of 3.40 Å and 3.48 Å for NTR– and TR-TLP, respectively (<u>Fig. S3</u>, <u>Table S1</u>). The 3D architecture of the treated and untreated spikes shows a similar average density. However, the NTR spike is slightly shorter and wider than the TR spike (Fig. 1B, E), and has an additional density at the base (Fig. 1C, F), consistent with the lectin domain of the VP4-C subunit previously described in these particles (Rodríguez et al., 2014; Shah et al., 2023). While the density of the VP2, VP6 and VP7 shells exhibit a homogeneous high local resolution, the spike shows a radial decrease in resolution as it extends away from the outer shell surface, with particularly low resolution in the body and head regions (<u>Fig. S3, S4</u>). This decrease may be related to the increased motion of the spikes as we move radially away from the particle. In addition, although the VP4-C lectin domain is detected, it is observed with low density (Fig. 1C), which has been associated to a flexible nature of its location (Rodríguez et al., 2014). These limitations hinder the construction of reliable VP4 atomic coordinates in the spike density and require further density improvement for accurate interpretation.

**Figure 1.**
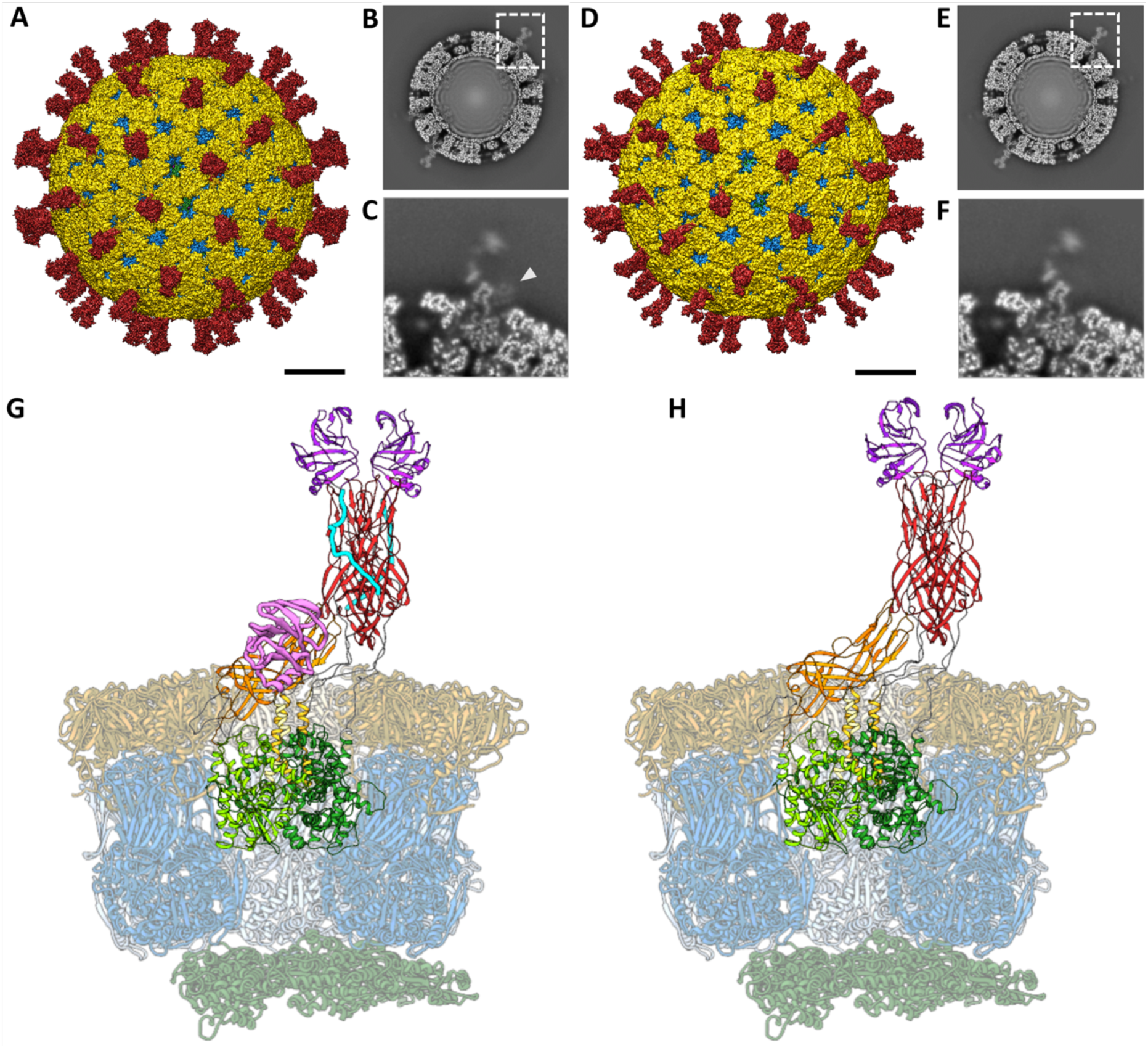
Analysis of the atomic structures of SA11 NTR– and TR-TLP by cryo-EM. (A, D) Cryo-EM 3DR of the NTR-(A) and TR-TLP (D) particles viewed along a 2-fold axis. The bar represents 100 Å. The surfaces are represented radially by colour coding: VP4 or VP8*/VP5* spikes (red), VP7 (yellow), VP6 (blue) and VP2 (green). The density is contoured to 2σ above the mean. (B, E) Cross-sections of the NTR-(B) and TR-TLP (E) maps, parallel to the central section, viewed along a 2-fold axis. (C, F) Close up view of the NTR (C) and TR (F) spikes indicated by a dashed square in B and E, respectively. The lectin domain of subunit VP4-C in the NTR spike is indicated by an arrowhead in C. (G, H) Atomic structure of the asymmetric subunit of NTR-TLP (G) and TR-TLP (H). VP2 chains are represented in dark green, VP6 in blue, and VP7 in yellow. The A, B and C chains of the VP4 or VP8*/VP5* spikes and their domains are represented as: foot (light green for the A chain, green for the B chain and dark green for C), body (red (A, B), orange (C)), head or lectin domains (purple (A, B), pink (C)). The binding loops between the different domains are represented in grey except for the α3-β13 loop, represented in cyan.

## Enhancement of spike resolution and interpretability

Technological and computational advances in recent years have made it possible to obtain structures of macromolecular complexes at atomic or quasi-atomic resolution using SPA techniques (Nakane et al., 2020; Yip et al., 2020). This approach relies on particle averaging and, while is especially powerful for resolving highly ordered and homogeneous structures, it is challenged when conformational or compositional structural heterogeneity is present in the sample. Conventionally, SPA pipelines rely on 2D and 3D classification to obtain particle subsets as homogeneous as possible. While this approach successfully addresses the compositional heterogeneity of the RV spike (partial occupancy) it is limited in addressing its continuous conformational flexibility (Rodríguez et al., 2014; Shah et al., 2023). In recent years, several new methods have been developed to refine structures with flexible and/or heterogenous components (Kimanius & Schwab, 2024). To improve the interpretability of our spike densities, we developed a pipeline that integrates localized reconstruction (Ilca et al., 2015), 3D classification (Zivanov et al., 2018), flexibility correction using optical flow (Kaur et al., 2021; Kazemi et al., 2021) and map enhancement and sharpening with LocSpiral (Kaur et al., 2021).

Icosahedral orientations determined for individual NTR and TR particles (n = 10,815 and 22,394, respectively) were used to subtract the contribution of the VP2/VP6/VP7 layers from the experimental images. Subsequently, subparticles at the spike locations (n = 648,900 and 1,343,640, respectively) were extracted using the localized reconstruction method described by Ilca et al. (Ilca et al., 2015) (<u>Fig. S5, S6</u>). The subparticles were sorted through 3D classification to separate capsid locations occupied by spikes from those not occupied. This resulted in 59% and 37% spike-occupied locations for NTR– and TR-TLP subparticles, respectively. The average density corresponding to the unoccupied subparticles was used to subtract the remaining VP6 and VP7 signal from the spike-occupied subparticles. Finally, 3D subclassification of spike conformations combined with optical flow was used to merge the slightly different subparticles into a unique conformation, thereby improving local resolution and signal-to-noise ratio (Kazemi et al., 2021). The amplitudes of the resulting maps were corrected by LocSpiral using a spiral phase transformation approach to enhance the interpretability of the obtanined 3DR (Kaur et al., 2021). The final density maps had the necessary quality and resolution required to model the atomic structure of both spikes. These densities, in conjunction with icosahedral 3DR of the whole NTR– and TR-TLP, were used to build and refine the polypeptide chains of VP2, VP4, VP6 and VP7 of the asymmetric subunits (<u>Table S1,</u> Fig. 1G, H).

## Spike rearrangement after activation

The structures of untreated and trypsin-treated TLP showed structural differences only in the spike (Fig. 1G, H <u>and</u> Fig 2), while the VP2, VP6 and VP7 layers are identical as previously described (Herrmann et al., 2021; Settembre et al., 2011; Shah et al., 2023). Both spikes exhibit the characteristic asymmetric arrangement, where the A and B subunits project outward from the particle as a dimeric body, while the C subunit lies prone in the stalk. As described previously (Rodríguez et al., 2014; Shah et al., 2023), the NTR spike displays the VP4-C lectin domain at the base of the stalk (Fig. 2A<u>, 4A pink</u>), which is absent after proteolytic processing. The density detected at the base of the stalk in the NTR spike allowed the modelling of the lectin domain in the chain C, from the residue T73 (Fig. 3A<u>, upper panels, green sphere;</u> Fig 4A<u>, left panel</u>) to residue N221 (Fig. 3A<u>, upper panels, blue sphere;</u> Fig 4A<u>, left panel</u>). While no density is observed for the loop linking K29 (<u>red sphere</u>) of the α domain to the T73 (<u>green</u> <u>sphere</u>), an empty density is present in the region between the lectin domain and the β-barrel of the C subunit (Fig. 3A<u>, upper panels, arrowhead)</u>. Although the map quality in this region is insufficient for building coordinates, its location is compatible with the 222-245 loop previously resolved by Shah et al. (Shah et al., 2023) that directly contacts the 414-420 beta hairpin from the VP4-C stalk. Since the biochemical analysis of the NTR-TLP (<u>Fig. S2A</u>) shows that the VP4 is not processed, the lack of density for the 29-73 and 245-253 regions (Fig. 3A<u>, upper panels, purple sphere</u>) is likely due to a high degree of flexibility in these loops.

**Figure 2.**
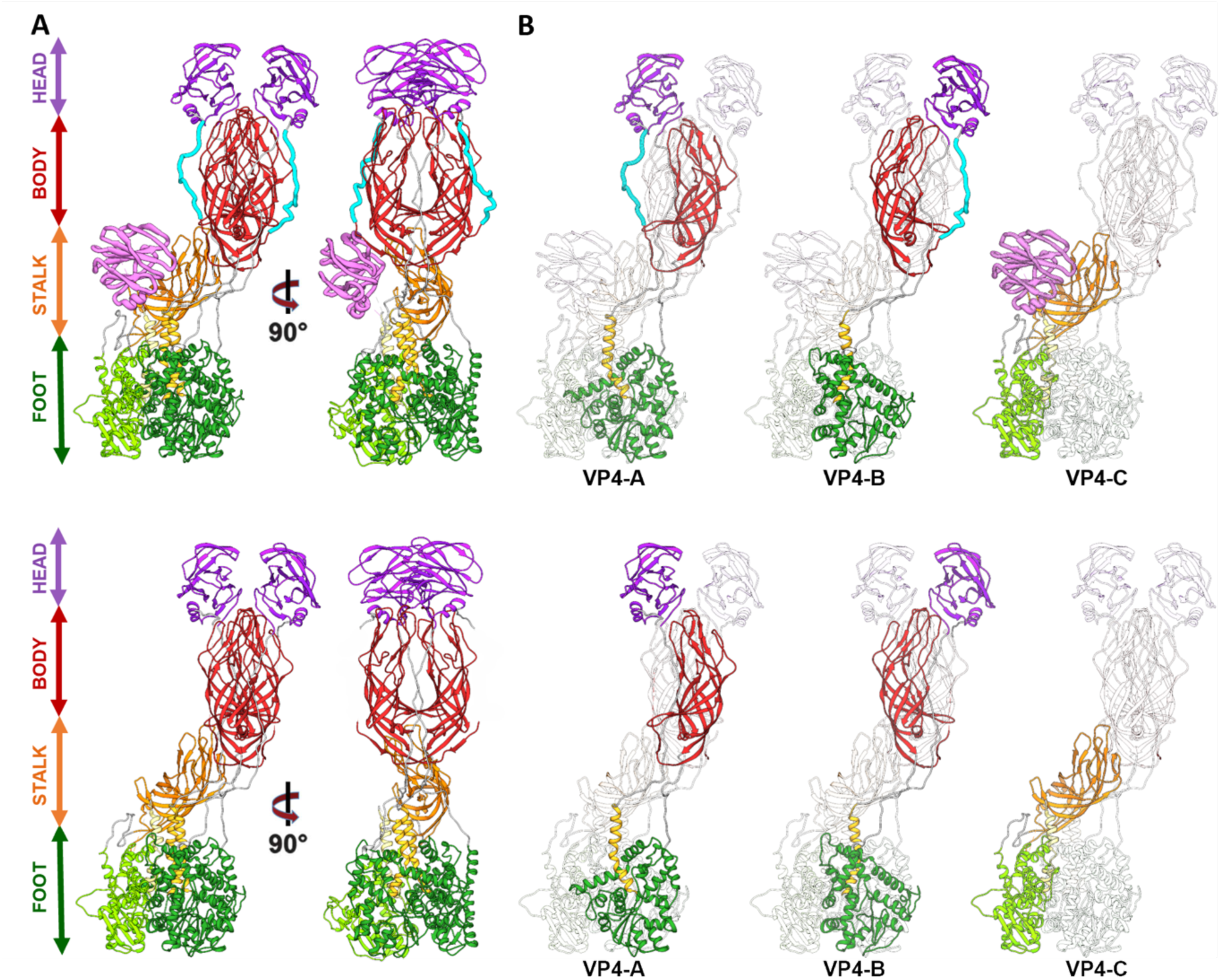
Atomic structure of the RV spike. (A) Atomic structure of the NTR (upper panel) and TR (lower panel) spike. Each domain is named and represented following the indicated colour pattern: foot (green), stem (orange), body (red), head (violet) and VP4-C lectin domain (pink, NTR spike). (B) Each of the VP4 subunits (A, B, C) of NTR-TLP (upper panel), and TR-TLP (lower panel) are shown highlighted and represented following the same colour pattern. In both panels, the binding loops between the different domains are represented in grey except for the α3-β13 loop, represented in cyan.

**Figure 3.**
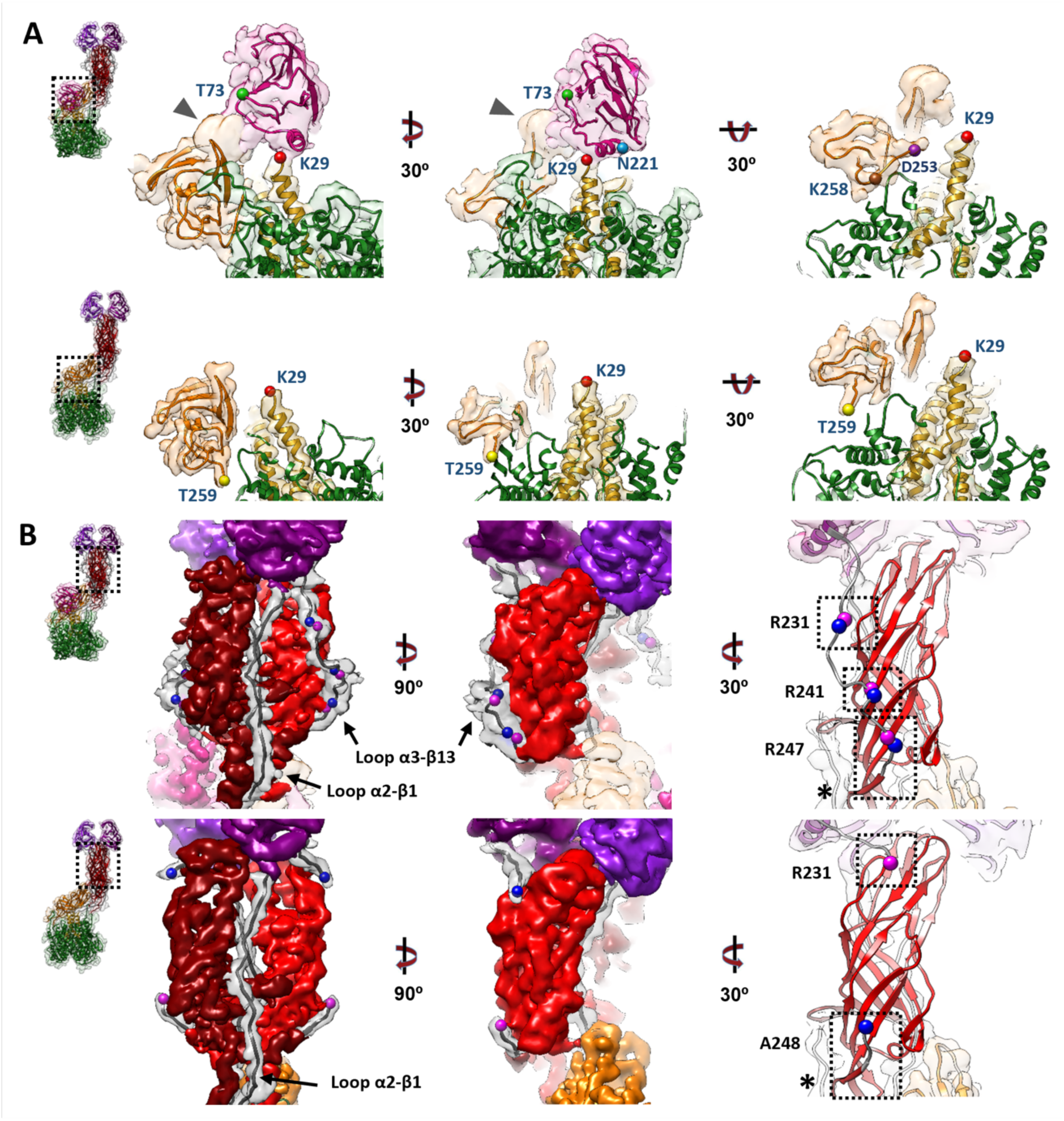
Analysis of the atomic structure of different domains of the NTR and TR spikes. A close-up view from the dashed boxes in the left panels of the atomic structures that have been fitted into their density maps. Within each panel (A and B) the upper row refers to the NTR spike and the lower row to the TR spike. (A) The stalk, lectin domain (VP4-C chain) and foot domains of the spikes are represented from different angles. Different residues are indicated as spheres: K29 (red), T73 (green), N221 (blue), K258 (brown), D253 (purple), T259 (yellow). (B) Side views of the body of the spike and the trypsinized (TR-VP4) or non-trypsinized (NTR-VP4) loops. Residues R231, R241, R247 are indicated in pink spheres and adjacent residues (N232, D242, A248) in blue. The α2-β1 and α3-β13 loops are shown. The molecular swapping observed in the body domain are indicated with an asterisk (⁕).

**Figure 4.**
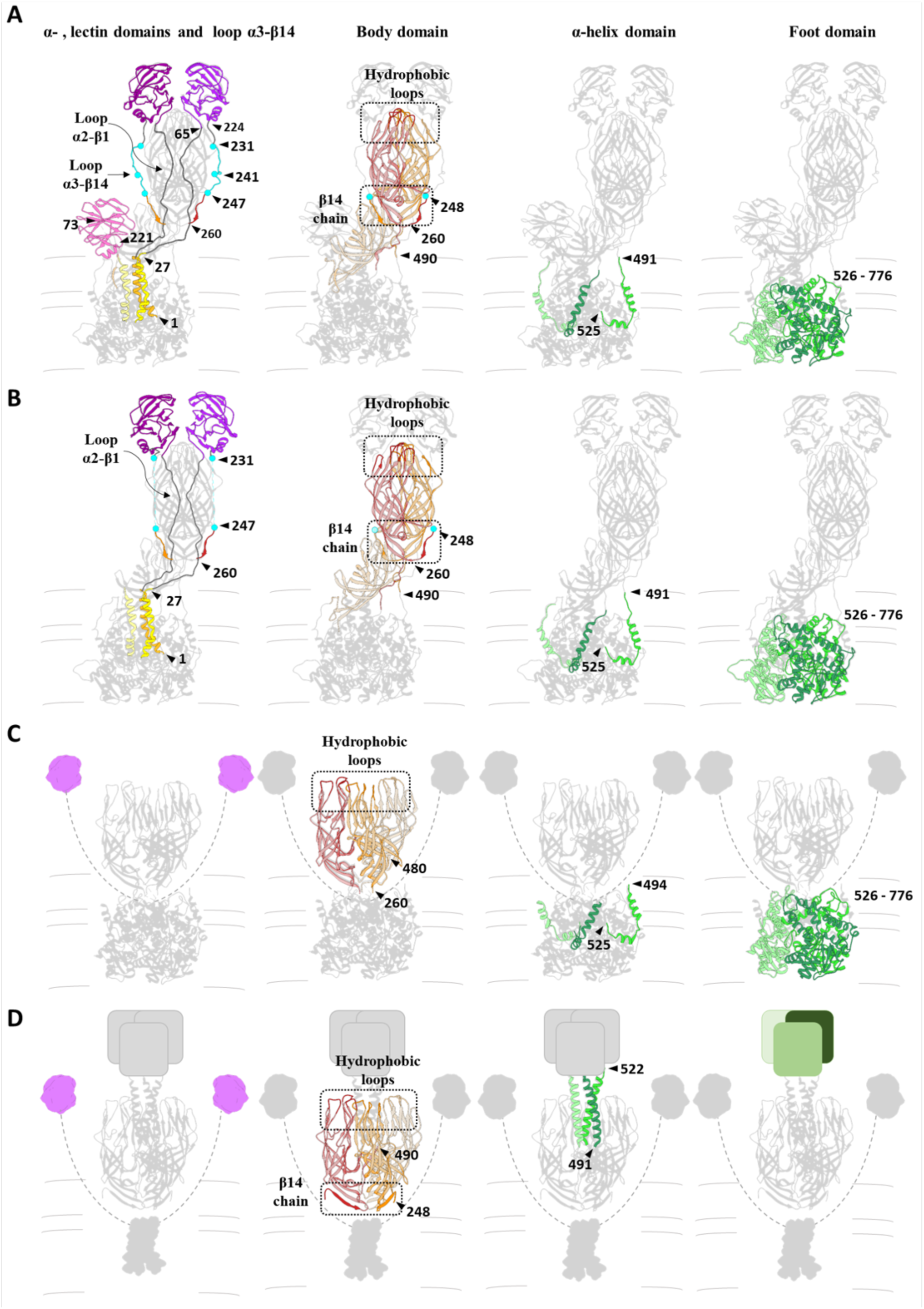
Analysis of the structural transitions of the loops and different domains of the RV spike during cell entry. Rearrangements of the spike proteins (VP4, VP8* and VP5*) during the transition from their inactive conformation, NTR (A), through their activated form, TR (B), and intermediate (C) to their inverted conformation. (D). The domains not observed in the atomic structures, in each case, are represented schematically (panel C: lectin domains; panel D: lectin, α-amino and foot domains). The different protein domains are coloured separately to illustrate their conformational change: α-amino domains: orange, yellow, gold; lectin domains: magenta, purple and pink; α3-β14 loops: blue; body domains: red, orange and salmon; α-coiled helix and foot domains: light green, green and dark green. Loops between the α-amino and lectin domains in panels C and D are represented by dashed lines. Arrows show different residues delimiting the different coloured domains in each panel. The panels corresponding to the body domain are shown with some transparency level and highlighting the hydrophobic loops and the β14 chain in a darker tone.

For the TR spike, the map allows for the modelling of the VP8*C chain from the first N-terminal residue to the K29 at the α2-β1 loop (Fig. 3A<u>, lower panel, red sphere;</u> Fig 4B<u>, left panel</u>). No electron density is observed between K29 and T259 (<u>yellow sphere</u>) of the stalk β-barrel domain. This region encompasses the lectin domain and the loops that connect it to rest of the spike structure. Biochemical analysis of TR-TLP (<u>Fig. S2B</u>) indicate complete proteolytic digestion of the spike. Considering the known trypsin processing targets at R231, R241, and R247 (C. Arias et al., 1996), the absence of density for the A248-K258 region could be attributed to either a high degree of flexibility in this region or an additional cleavage after K258 for the C subunit (Fig. 3A<u>, brown sphere</u>). The first defined residue in the stalk β-barrel of the previously resolved structure for the RRV strain TR-TLP is Glu264, suggesting cleavage of the C-terminal to either K258 or K263 (Settembre et al., 2011). Previous analyses partially evidence a cleavage at K258 (Crawford et al., 2001; P. R. Dormitzer et al., 2001). Our structure, as well as the structure of VP4/VP7 recoated TLP (Herrmann et al., 2021), support this scission. On the other hand, the structure of the inverted conformation of the spike solved by Herrmann et al. shows the presence of residues 248-258 (α3-β14 loop) in the three VP5* subunits, which implies that under these conditions the chain C is not processed at K258.

The 3DR of the NTR spike allows for the atomic modeling of the entire polypeptide chain of the A and B subunits, including the two long loops that connect the head lectin domains to the rest of the molecule: the internal α2-β1 loop, which connects the α domain with the lectin domains, and the α3-β13 external loop, which binds them to the body of the spike (Fig. 2A, B, cyan; Fig. 3B, upper panels; Fig. 4A, left panel, <u>Video S1</u>). Complete cleavage of the α3-β13 loop is evidenced as an abrupt loss of signal in the density map after R231 and before A248 (Fig. 3B<u>, lower panels;</u> Fig. 4B<u>, left panel</u>, <u>Video S1</u>). After cleavage, the 225-231 amino region of the cleaved α3-β13 loop retracts and establishes new contacts with the residues 443-448 in the β-barrel domain of the opposite chain (Fig. 3B<u>, right panels;</u> Fig. 4B<u>, left panel</u>, <u>Video S1</u>).

The carboxy terminal region maintains its interaction with the β14-sheet present in the unprocessed spike and contributes to the stability of the spike structure. (Fig. 3A, <u>right panel, asterisk;</u> Fig. 4A, B <u>central panels</u>). The head lectin domains maintain a similar structure before and after cleavage, with their glycan binding sites equally accessible to their host cell targets (<u>Fig. S7</u>), which correlates with the binding capacity of the virions to the target cell independently of the proteolytic state of their spikes (Clark et al., 1981).

The structure of the NTR spike reveals that α2-β1 and α3-β13 loops that connect the lectin domains to the preceding α domain and the following β-barrel domain embrace the body of the spike and firmly anchor the head on top of the spike. Actually, these loops maintain the NTR spike in a conformation slightly more compact than that of to the TR spike. After trypsin activation, the suppression of the physical constrains imposed by the α3-β13 loop is reflected as an expansion of the spike domains orthogonally to the capsid surface. While the foot and stalk domains do not suffer significant movements between their α-carbon chains, with a root-mean-square deviation (rmsd) of 1.1 Å and 1.5 Å, respectively; the rmsd of the body and head domains are 3.3 Å and 5.7 Å, respectively. In this activated conformation the head domains are only anchored to the foot by the α2-β1 loop, making them competent for the lateral movement of the lectin domains that will initiate the structural transitions that finalize with the host cell membrane penetration (<u>Fig. S1C</u>).

## Discussion

Proteolytic cleavage of the spike protein is known to be essential for RV infection, but the structural details of how cleavage within VP4 transitions the RV particle from its non-infectious immature state to the infectious mature state have remained elusive. The intrinsic flexibility of the spike structure and the low occupancy of spikes on the virion have significantly hampered efforts to determine the structural determinants behind this critical molecular mechanism. Here, by employing a combination of advanced computational methods (Ilca et al., 2015; Kaur et al., 2021; Kazemi et al., 2021), we have successfully overcome these limitations to construct an atomic model of the non-processed and processed NTR spike. Together with the contributions of others, this structure completes the comprehensive model of the conformational pathway that the RV spike follows from its assembly in the ER to membrane penetration in a new host cell (Fig. 4<u>, 5</u>).

The lectin domain in the VP4-C of the mature NTR spike, previously described (Rodríguez et al., 2014; Shah et al., 2023) (Fig. 4<u>, 5</u>), is positioned close to the capsid, making it intrinsically less accessible to cell surface receptors. The VP7 layer further restricts access to site II in the galectin-like fold structure (<u>Fig. S7</u>). Thus, it is highly likely that the ability of NTR-TLP to bind the cell membrane before spike activation by proteolysis (C. Arias et al., 1996; Gilbert & Greenberg, 1998; Graham & Estes, 1980; Rodríguez et al., 2014) is mediated by the head lectin domains, although this binding cannot initiate a productive infection. Moreover, VP4-C lectin domain is cleaved following trypsin proteolysis in the TR spike, suggesting that its glycan-binding function may be dispensable during subsequent stages of cell entry. This domain may serve an unknown specific function in the NTR spike, or could be a vestigial component of the spike structure, which has evolved from three identical subunits into a partially trimeric (foot) and partially dimeric (body-head) structure that must transform into a fully trimeric configuration to engage with and penetrate the membrane.

Spike proteolytic activation is mediated by the cleavage of the α3-β13 loop in three target residues that have different susceptibilities to trypsin, R241 > R231 > R247 (C. Arias et al., 1996). The central region of the α3-β13 loop, including the 232-242 region, is more distant from the spike body and does not establish direct interactions with the β-barrels. In contrast, the preceding and following regions, which contain residues R231 and R247, are contacting the adjacent β-barrel. This structural arrangement clarifies the molecular basis of the differential susceptibility to trypsin: the central region of the loop, being more exposed and less constrained, is more accessible to the protease, making it more susceptible to cleavage.

The determination of the structure of the α3-β14 loop in the NTR spike allows to understand why the activation by proteolysis is necessary for infectivity. In non-activated virus the spikes have their lectin head domains covalently linked to the foot and the body domains through the internal α2-β1 and external α3-β14 loops (Fig. 5). Furthermore, the external α3-β13 outer loops embrace the opposite β-barrel in the body of the spike, mediating a molecular swap that enhances the stability of this structure. In his conformation, the presence of the α3-β14 loop prevents the displacement of these lectin domains, acting as a “lock” preventing further conformational changes. The breaking of the covalent bonding and the loss of the segment 232-247 releases the structure from his locked state and allows for the lateral displacement of the VP8* lectin domains, which initiates the conformational transitions that mediate virus entry. It is noteworthy that when the R247 trypsin site in the α3-β14 loop of the rotavirus strain KU is mutated to a furin site, endogenous furin-like proteases cleave the mutant VP4 intracellularly after TLP assembly, and results in the formation of a mutant virus that is incapable of plaque formation (Komoto et al., 2011). This unexpected outcome has been attributed to an inefficient process of virus release (Komoto et al., 2011). After its synthesis in the endoplasmic reticulum (ER), furin travels through the Golgi apparatus and reaches the trans-Golgi network (TGN), also cycling between the TGN, cell surface, and endosomes. This makes it likely that immature spikes on nascent TLPs within the ER are prematurely activated by furin, leading to their interaction with intracellular membranes and effectively trapping the nascent viruses within the cell due to the presence of cleaved spikes. Different human (Wa, DS-1) and animal (SA11, RRV, OSU) RVA species shows a clear conservation of both the canonical sites for trypsin digestion, R231, R241, and R247, as well as the K29 and K258 sites (<u>Fig. S8</u>). Only in the DS-1 strain is this conservation not observed at R231, although it is maintained throughout the rest of its sequence. Extending this comparison to other rotavirus groups (B-D, F-I) (<u>Fig. S8</u>), all of them have potential trypsin sites at or near residue 29. Except for RVA, the other groups have at least two trypsin sites in the α2-β1 loop and some of them, such as B, G, H and I, have multibasic cleavage sites, which present between 5 and 9 sites susceptible to trypsin proteolysis. Similarly, the α3-β14 loop has multibasic cleavage regions with 4 to 9 potential trypsin sites in all RV groups. The presence of additional monobasic residues in both, α2-β1 and α3-β14, loops in some groups of RV could serve as a mechanism to ensure the virion activation in the gastrointestinal tract, where newly assembled particles are released.

**Figure 5.**
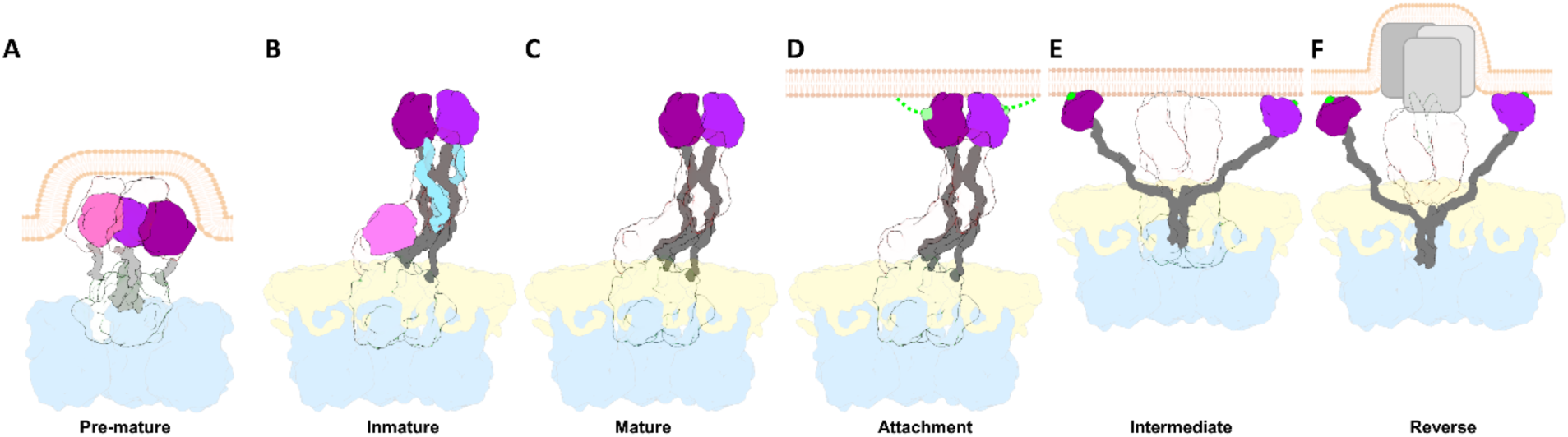
Structural transitions of the rotavirus spike. Schematic representation of the proposed conformational changes of the RV spike during its morphogenesis, maturation and activation. Proteins are coloured as indicated: VP6 in blue, VP7 in yellow, VP4/VP5* foot and β-barrel transparent, VP4/VP8* lectin domain A, B and C in magenta, purple and pink, respectively, and loops in grey. (A) During the late stages of ER morphogenesis, full-length VP4 monomers fold into a pre-mature three-fold symmetric structure in the eDLP. It has been proposed that the transition from this pre-mature conformation to the immature conformation is one of the driving forces for the transition of the eDLP to TLP. (B) In the immature TLP, the three unprocessed VP4 subunits are assembled as an asymmetric trimer. Two VP4 subunits. chains A and B, form the body and head of the spike joined by two internal (gray) and two external loops (light blue). The VP4-C chain folds to form the stalk of the spike to which it contributes with the β-barrel and the lectin domain. (C) Trypsin activation splits VP4 into VP5* and VP8* products in the mature spike. Trypsin proteolysis occurs at the external α3-β14 loop in three residues, R231, R241 and R247, leading to the loss of the segment 232-247 (loss of light blue loops) in the VP4-A and –B chains, and the loss of the lectin domain in the VP4-C chain (pink). (D) The activated spike is able to bind to the host cell through the interaction of VP8* lectin domains with surface glycans (light green) that result in the attachment of the virus to the cell membrane. (E) This interaction precedes the conformational change in which the lectin domains separate from the main body of the spike, exposing the hydrophobic loops of the ß-barrel domains of chains A and B. This conformational change continues with the extension of the ß-barrel domain of subunit C and ends in the intermediate conformation in which the ß-barrel domains of the three subunits adopt a symmetrical trimeric structure with the three hydrophobic loops inserted into the target membrane. F) After this interaction a further conformational transition results in the inverted conformation, in which the loops and the foot domains of the spike are insert into the membrane which is thought to provoke membrane distortions which culminate in its rupture and in the release of the DLP into the cytosol.

The precise timing and location of the spike conformational transitions are highly regulated to ensure the success of the viral infectious cycle. The fact that the non-enveloped rotavirus virion assembles within the ER is particularly remarkable and likely relates to the intricate conformational dynamics of the virion spike, an extreme example of polymorphism for the VP4 polypeptide. Previous studies (Shah et al., 2023) have demonstrated that VP4 initially assembles as a membrane-interacting, 3-fold symmetric pre-mature structure, stabilized by charge-based and hydrophobic interactions between the lectin domains and the side of the β-barrel. The transition from this form to an immature state involves a simultaneous extension of VP4 into an upright, elongated ‘dimeric’ asymmetric conformation, accompanied by the addition of the VP7 layer and the loss of the transient envelope. Stabilization of the immature spike occurs through increased lateral interactions between the elongated A and B subunits, while the C-subunit head domain remains positioned at the base of the uncleaved VP4 spike. This conformation is maintained within the ER in nascent TLPs and continues to exist in the extracellular immature TLP (Rodríguez et al., 2014; Shah et al., 2023). To ensure successful infection, the spike structure is ‘locked’ by the α2-β1 and α3-β13 loops, which anchor the lectin domain that conceal the hydrophobic loops until the moment of infection. To avoid premature activation, RV has evolved the structure and sequence of these loops to ensure activation occurs only when they encounter trypsin-like proteases in the intestine, preventing early conformational changes and exposure of the hydrophobic loops.

## Materials and Methods

### Virus production, purification and titration

In this study, the C4111 clone of the SA11 simian rotavirus A strain (RVA/Simian-tc/ESP/SA11-C411/2009/G3P[2]) (Rodríguez et al., 2014) and the African green monkey kidney epithelial cell line MA104 (ECACC 85102918) were used to obtain rotavirus particles. Cells were grown in MEM medium supplemented with 2 mM glutamine, 50 µg/ml gentamicin and 10% foetal serum, maintained at 37°C with 5% C02 and 95% humidity and used between passages 7 and 24. The purification of TLP produced in the presence or in the absence of trypsin was performed as previously described (Rodríguez et al., 2014). Viral infectivity was determined by plaque assays as previously described (Rodríguez et al., 2014).

### Electron microscopy and image processing

For cryo-EM analysis, copper-rhodium or gold Quantifoil R2/2 holey grids were ionized by ion discharge (Q150T, Quorum). Samples were applied, blotted, and plunged into liquid ethane using a Leica EM GP2 automatic cryo-fixation unit. Grids were analyzed under low-dose conditions in different Titan Krios microscopes (Thermofisher Scientific) operated at 300 kV using different direct detectors in linear mode. In all cases, data collection was performed with the automatic acquisition program EPU (Thermofisher Scientific) for single particle analysis. The acquisition conditions for each of the samples are indicated in Table 1.

Image processing operations were performed using Xmipp (de la Rosa-Trevín et al., 2016) and RELION (Scheres, 2012; Zivanov et al., 2018) integrated into the Scipion (de la Rosa-Trevín et al., 2016) platform and graphic representations were produced by UCSF Chimera (Pettersen et al., 2004). MotionCor2 (Zheng et al., 2017) routine was used to align, dose correct and average frames of the acquired movies. The contrast transfer function (CTF) and defocus were later estimate with CTFFIND4 (Rohou & Grigorieff, 2015) and Xmipp automatic picking routine was used to select particles (indicated in Table 1). Images were 2D classified using the corresponding Relion routine and the selected particles were analyzed using the 3D classification routine. Given the size of the particles, and to obtain greater computational efficiency, during the 2D and 3D classification steps, the particles were subsampled three times with respect to the original sampling frequency. We imposed icosahedral symmetry for alignment and reconstruction. The corresponding Relion auto-refinement routine was used with the original, non-downsampled, particles to obtain a 3DR and resolution was assessed by gold standard Fourier Shell Correlation (FSC) (van Heel, 1984) between two independently processed half datasets (Scheres & Chen, 2012). The resolution of each reconstruction (Table 1) was determined applying a correlation limit of 0.143. Additionally, the local resolution of each map was determined with the MonoRes routine (De la Rosa-Trevín et al., 2013; Vilas et al., 2018). Next, a correction of the amplitudes at high frequencies was performed using the RELION Postprocessing routine, which automatically determines a B-factor to apply to the cryo-EM map weighted by the curve of the FSC. The values obtained from B-factor for each map are found in Table 1. Additionally, amplitudes at high frequencies were corrected using the Xmipp LocalDeblur routine (Ramírez-Aportela et al., 2021) which perform a correction based on the local resolution.

The asymmetric subunits of rotavirus particles were extracted using the UCSF Chimera program (Pettersen et al., 2004) or the Xmipp program (ExtractUnitCell) (De la Rosa-Trevín et al., 2013; Sorzano et al., 2013).

### Determination of the spike 3D maps

The localized reconstruction method (Ilca et al., 2015) was used for both 3D maps, NTR– and TR-TLP. This method allows the generation of 3DR of VP4 spike regions that present higher flexibility, different symmetry, and degrees of occupancy. These regions, hereinafter called subparticles, are studied as isolated particles, thus facilitating their classification and subsequent refinement and 3DR steps (<u>Fig. S5</u>). The signal contribution of the VP2, VP6 and VP7 layers was subtracted from the original TLP 3DR using the density map of the corresponding TLP and a mask that encompasses these three layers. Subsequently, considering the position of the spikes in the final 3DR and its icosahedral symmetry, the 2D projections of each spike were extracted from the difference images calculated previously. Subsequent 3D classification of these VP4 subparticles of the TLP spikes generated in the absence (NTR spikes) and presence of trypsin (TR spikes) revealed in both cases a class with a density corresponding to VP4 and another class in which the density of VP4 was absent. The surrounding VP7 and VP6 density was subtracted using the non-spike density class from the map with the density of VP4. In the case of NTR spikes, a second 3D classification round was performed to identify subparticles with a density associated with the VP4-C lectin domain. Next, 3D maps of both NTR and TR spikes were reconstructed. Additionally, the EnRICH (Resolution Improvement in Conformational Heterogeneity maps) method was implemented for NTR and TR spike reconstructions as previously described (Kazemi et al., 2021). This method transforms the particles from their initial conformation to an established reference conformation. For this, the following successive steps were carried out: 1) 3D classification into 6 classes to obtain different conformations of the map and particles associated with them; 2) 3D refinement of each class to obtain the precise angular assignments of each particle; 3) translocation of the particles with different conformations to a reference one through an optical flow process and 4) alignment and 3D reconstruction of the set of all the particles with the new conformation with better resolution, contrast and signal-to-noise ratio As a final step, the LocSpiral algorithm (Kaur et al., 2021) was applied to improve the interpretability of the 3DR.

### Structure modelling, refinement and validation

The asymmetric unit of the NTR– and TR-TLP SA11 rotavirus strain under study consist of the trimetric VP4 spike protein and 4 + 1/3 trimers of each VP6 and VP7. We initially generated a homology atomic model for each polypeptide chain (VP2, VP4, VP6 and VP7) that forms the asymmetric subunit of the capsid using the published atomic structures of the rotavirus RRV TLP (PDB: 4V7Q (Settembre et al., 2011) and 6WXE (Herrmann et al., 2021). The reference PDB was adjusted in the density map and the chains were generated by homology modeling with the Modeller program (Šali & Blundell, 1993) within UCSF Chimera (Pettersen et al., 2004) by pairwise alignment of sequences. The chains were adjusted as a rigid body in the density map with the UCSF Chimera program (Pettersen et al., 2004). Coot program (Emsley & Cowtan, 2004) was used to manually trace or adjust the polypeptide chains in the map. In each case, the map with the amplitude correction that best allowed us to carry out successive atomic modeling processes was selected.

Finally, iterative rounds of refinement and validation of the model were carried out using PHENIX programs (Adams et al., 2010) (Table 1). In each case, the quality of the model and the correct geometry of the atomic structure was validated through the statistics determined by Molprobity (V. B. Chen et al., 2010). We used standard stereochemical and B-factor restraints, as well as Ramachandran, rotamer, and secondary structure restraints. Residues included in the models and model statistics are summarized in Table 1. Molecular graphics and analyses were performed with UCSF Chimera (1.18), developed by the Resource for Biocomputing, Visualization, and Informatics at the University of California, San Francisco, with support from NIH P41-GM103311 (Pettersen et al., 2004).

### Multiple sequence alignment analysis

Multiple sequence alignments of VP4 protein sequences of representative strains of the RVA specie (Wa, DS-1, OSU, RRV, SA11) and the reference strains for the different RV species were performed using the Clustal Omega server (Goujon et al., 2010; Sievers et al., 2011) and analyzed using the ESPript server (ESPript – https://espript.ibcp.fr; (Robert & Gouet, 2014)). We retrieved rotavirus sequences from GenBank (Benson et al., 2018). The accession numbers of the VP4 genomic segments of the different strains used are the following:

| **RV specie** | **Strain** | **Accession number** |
| --- | --- | --- |
| A | SA-C4111 | AIC34758.1 |
| A | DS-1 | AEG25325.1 |
| A | Wa | P11193.3 |
| A | OSU_C5111 | AIC34770.1 |
| A | RRV | P12473.2 |
| B | Bang373 | YP_008126845.1 |
| C | Bristol | CAA55958.1 |
| D | 05V0049 | YP_003896048.1 |
| F | 03V0568 | YP_008145319.1 |
| G | 03V0567 | YP_008136232.1 |
| H | J19 | YP_392492.1 |
| I | KE135 | YP_009130677.1 |

### Accession codes

Structures described in this manuscript have been deposited in Protein Data Bank under accession code 8OLB and 8OLC for the NTR– and TR-TLP structures, respectively. The cryo-EM data has been deposited and issued the code EMD-16954 and EMD-16955, for the NTR– and TR-TLP structures, respectively.

The NTR– and TR-spikes structures have been deposited in Protein Data Bank under accession code 8OLE and 8QTZ, respectively. The cryo-EM data has been deposited and issued the code EMD-16956 and EMD-18655, in each case.

|  | **Protein Data Bank (PDB)**<br>**accession code** | **Electron Microscopy Data**<br>**Bank (EMDB) code** |
| --- | --- | --- |
| **NTR-TLP** | 8OLB | EMD-16954 |
| **NTR spike** | 8OLE | EMD-16956 |
| **TR-TLP** | 8OLC | EMD-16955 |
| **TR spike** | 8QTZ | EMD-18655 |

## Acknowledgements

This work was supported by grants of the Instituto de Salud Carlos III (PI20CIII-00014) to DL and of the Spanish Ministerio de Ciencia e Innovación (PID2022-137548OB-I00 funded by MCIN/AEI/10.13039/501100011033/) and by ERDF A way of making Europe to JV.

## Author contributions

Conceptualization, DAC, JMR and DL; Methodology, all authors; Validation, CPM and DAC.; Formal analysis, all authors; Investigation, all authors; Writing-Original Draft, DAC, JMR and DL; Funding Acquisition, JV, JMR and DL.

## Supplementary Information

**Figure S1.**
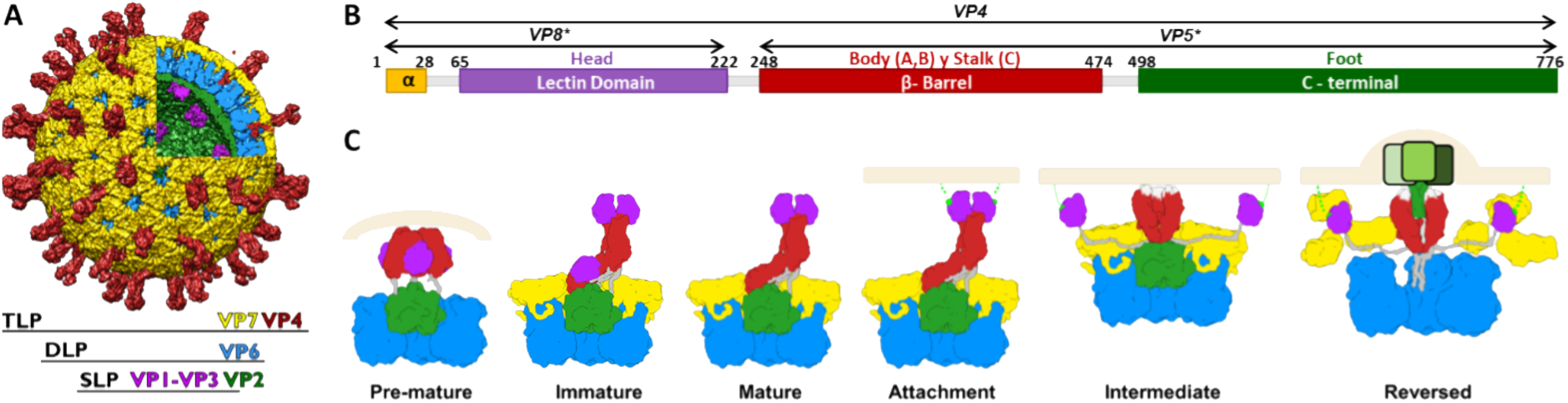
Rotavirus structure and structural transitions of the RV spike during the viral cycle. (A). Representation of the RVA viral particle. The different protein layers ensembled and their structural proteins are indicated with the colour code. (B). Schematic representation of the structure and domain organization of VP4. The atomic structure (PDB: 4V7Q) is represented showing each of the monomers that forms the RV spike (VPA-A, VP4-B, VP4-C) and the names of its domains. The panel represent the primary structure of the spike indicating the different domains: α (yellow), lectin (magenta), β barrel (red), and C-terminal (green) domains. The VP4 proteolytic products (VP5* and VP8*) and domains are labelled. Residues delimiting domains and trypsin cleavage sites are indicated. (C). Structural transition of the rotavirus spike during the infectious cycle. Proteins are coloured as indicated: VP6 in blue, VP7 in yellow, VP4/VP5* foot in green, VP4/VP5* β-barrel in red, VP4/VP8* lectin domain in magenta, and loops in grey. During the last stages of the morphogenesis in the endosome, the full-length VP4 monomers in the pre-mature TLP form a flexible 3-fold symmetry structure which carry out a conformational change into the upright structure found in the immature TLP. In this immature spike, two VP4 subunits assemble forming the body and head of the spike (A and B chains) joined by loops (gray). The third VP4 subunit folds to for the stalk with a β-barrel and a lectin domain (C chain). Trypsin proteolysis cleaves the VP4 chains into VP5* and VP8* subproducts. The trypsinization of the α3-β14 loop in three residues, R231, R241 and R247, leads to the loss of the segment 232-247 in the VP4A-B chains and the loss of the lectin domain in VP4C in the mature spike. The activated spike a to the host cell through the interaction of VP8* lectin domains (attachment) with surface glycans (light green) of the cell membrane. This interaction precedes the conformational change in which the lectin domains separate and expose the hydrophobic loops resulting in an intermediate conformation in which these loops interact and insert themselves into the membrane, causing its distortion, rupture and DLP releasing in the cytosol (reversed conformation).

**Figure S2.**
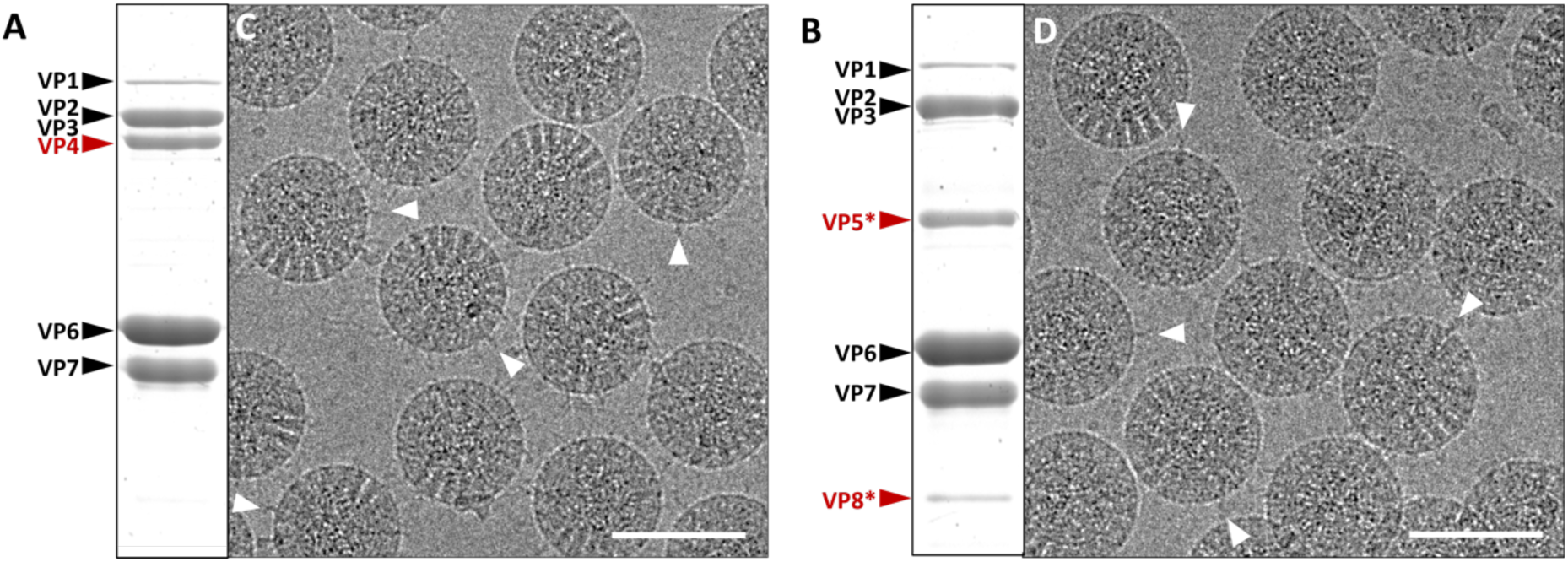
Purified RV SA11 NTR– and TR-TLP analyzed by SDS-PAGE and cryo-electron microscopy. (A, B) Coomassie blue-stained SDS-PAGE of purified TLP, cultured in the absence (A) or presence (B) of trypsin. The positions of RV structural proteins (VPs) are indicated. (C, D) Cryo-electron micrographs of NTR-(C) and TR-TLP (D). The position of some spikes projected from the surface of the particles is indicated with white arrowheads. The bar represents 100 nm.

**Figure S3.**
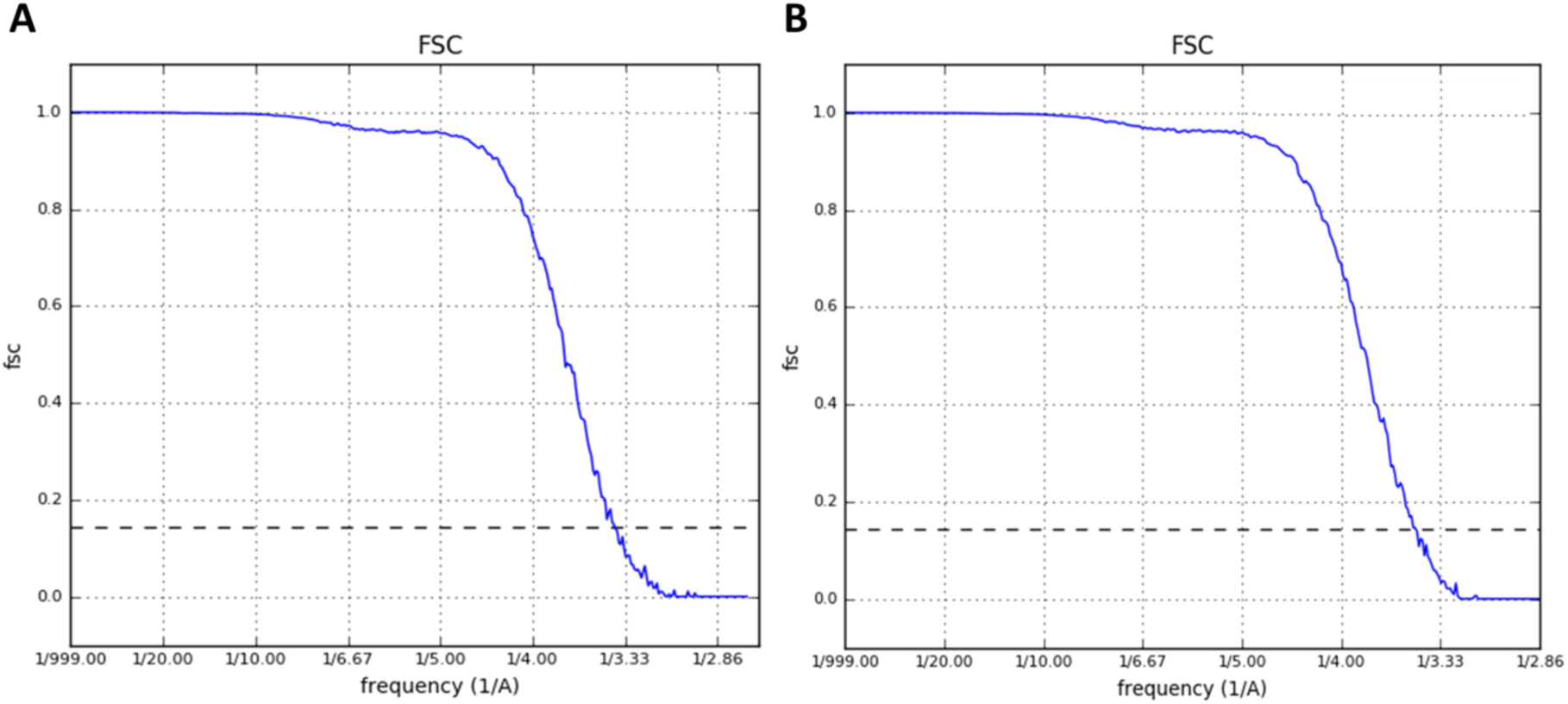
Resolution of the atomic models of the NTR-TLP and TR-TLP. The resolution values of the NTR-TLP, 3.40 Å, and TR-TLP, 3.48 Å, are based on the FSC criterion at 0.143.

**Figure S4.**
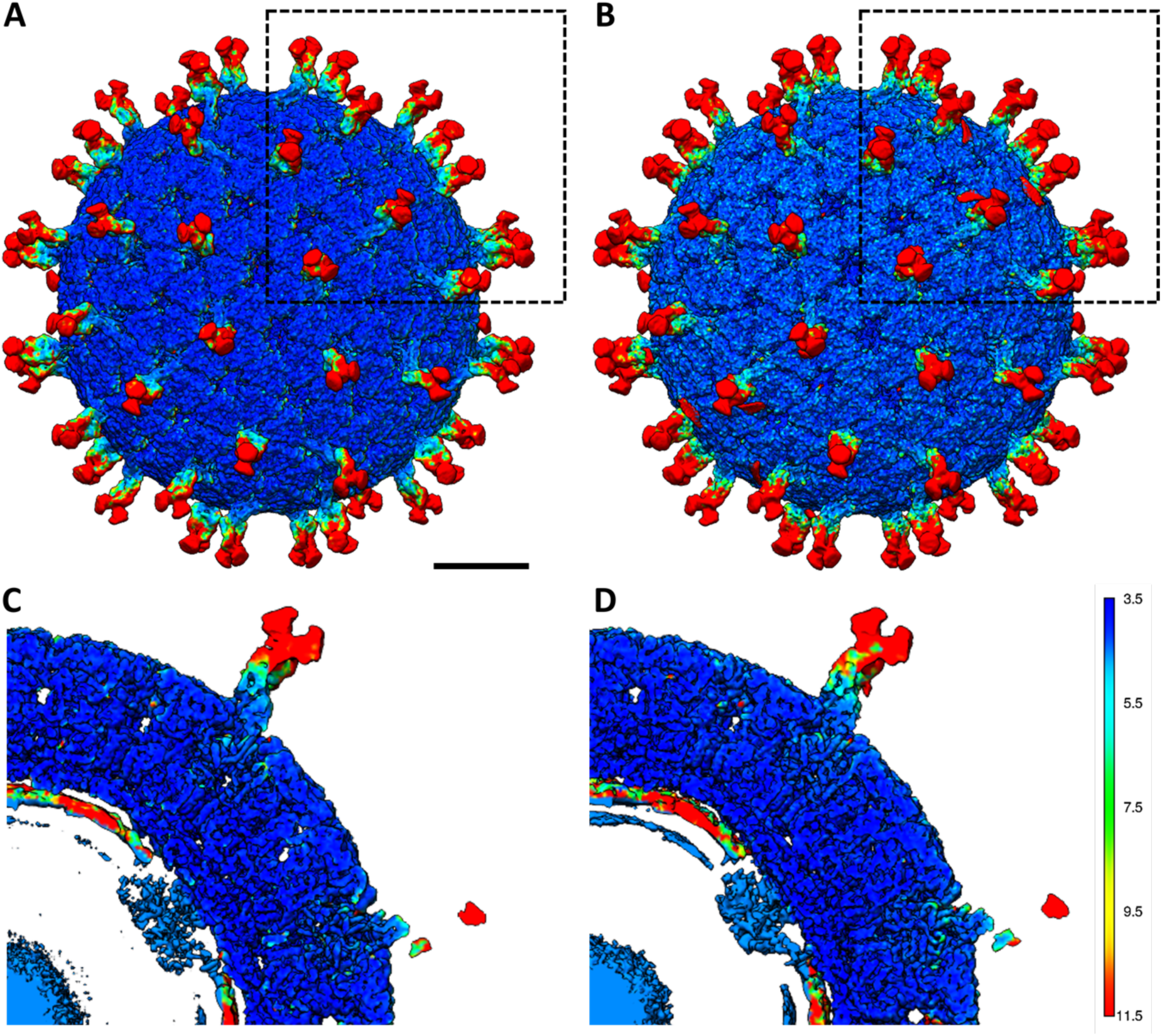
Analysis of the local resolution of the 3D maps obtained from the NTR– and TR-TLP. (A-B) Representation of the 3D maps of the NTR-(A) and TR-TLP (B) particle viewed along the icosahedral axis of symmetry 2. The densities observed in panels C and D are indicated with a dashed square. (C-D) Close view of NTR-(C) and TR-TLP (D) cross sections of each 3D map. The sections are parallel but offset 14.7Å from the central section of the maps. The surfaces are coloured according to the local resolution calculated for each 3DR. The colour code is shown with the corresponding resolutions in Å. Densities are contoured at 2σ above the mean. Scale bar represents 100 Å.

**Figure S5.**
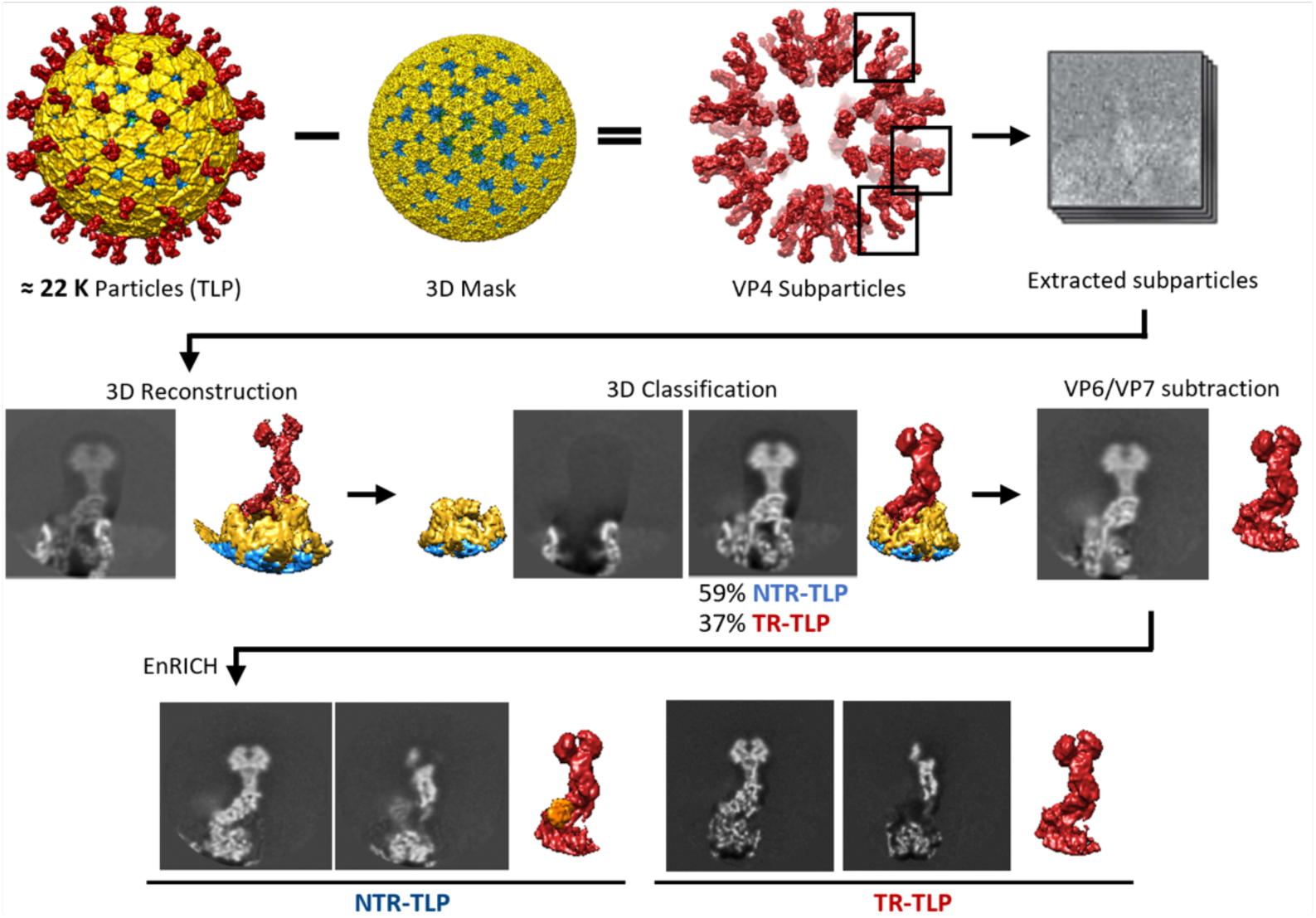
Digital processing performed to obtain the 3DR of the spikes. Firstly, we used the localized reconstruction method (Ilca et al., 2015): the VP2, VP6 and VP7 layers were subtracted using the corresponding TLP maps and a mask that encompasses these three layers. Subsequently, the spikes from all positions were extracted from the calculated difference images and treated as individual particles for their 3D classification, refinement and reconstruction. A 3D classification separated the positions occupied and not occupied by spikes, with an occupancy level of 59 and 37% for NTR– and TR-TLP, in each case. The unoccupied 3DR were used to subtract the VP6 and VP7 signal from the spike-occupied subparticles. Finally, the EnRICH method was applied (Kazemi et al., 2021) to obtain aligned subparticles whose 3D reconstructions showed a significant increase in their local resolutions, contrast and signal-to-noise ratio in the stalk, body, and head regions for both spikes.

**Figure S6.**
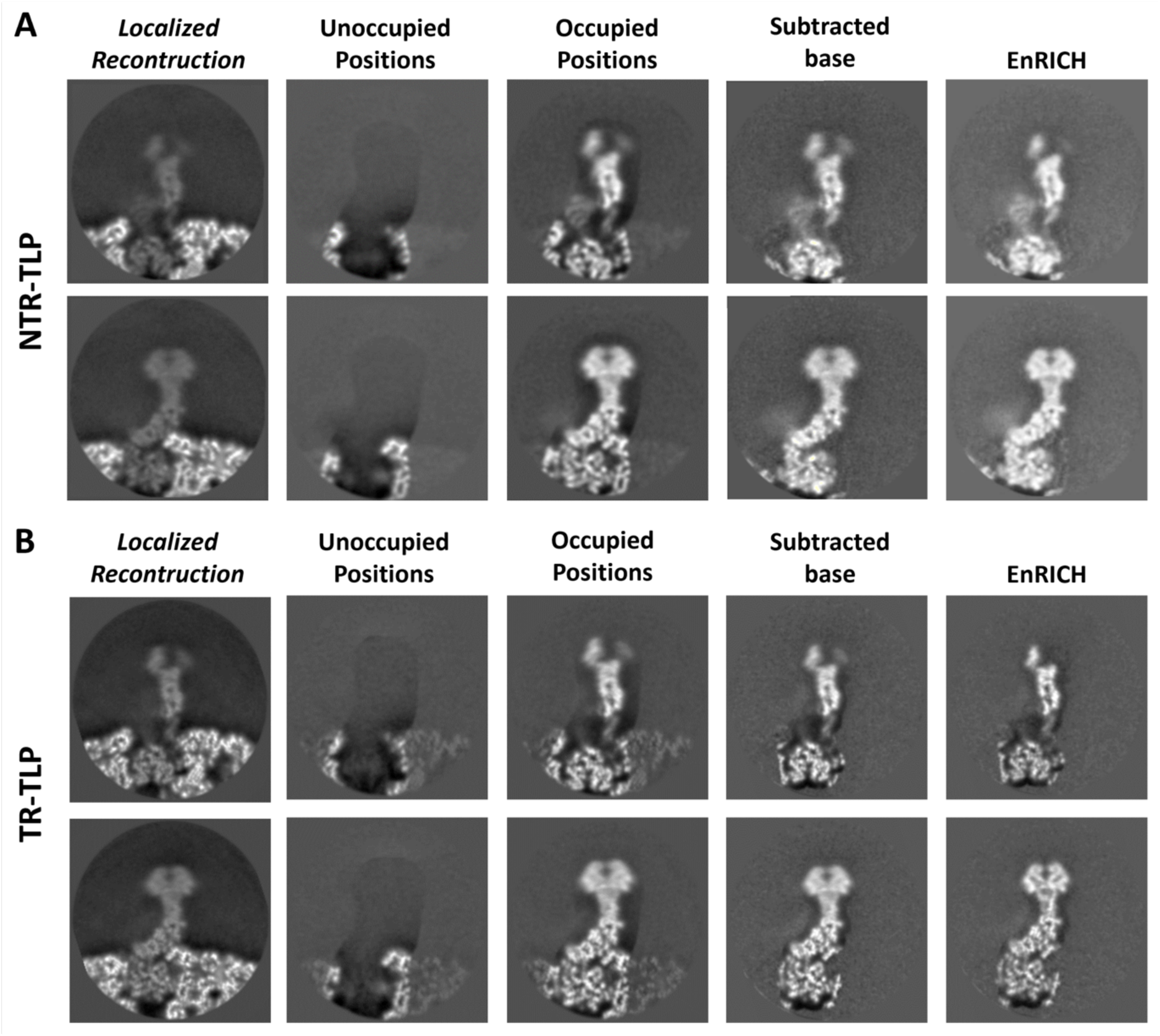
Analysis of the local resolution of the NTR– and TR-spikes at the different stages of the subparticle refinement. 1.34 Å thick cross sections of the maps obtained at the different stages of refinement of the VP4 NTR and TR subparticles. The panels show sections of each 3D map parallel to the central section of the maps and offset by 14.7Å (top panels) and 6.7Å (bottom panels).

**Figure S7.**
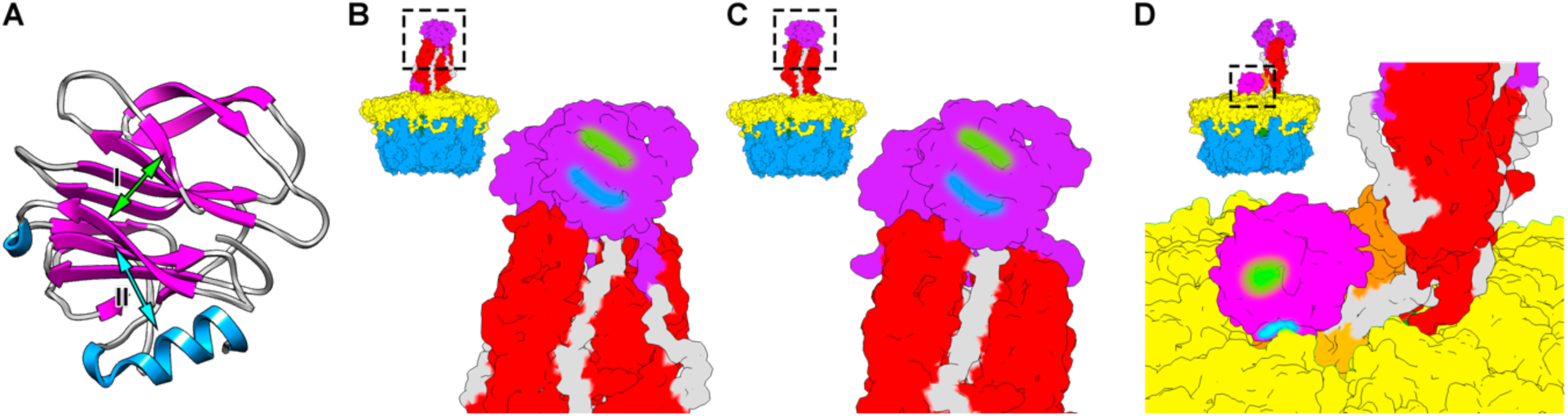
Accessibility of the glycan binding sites of the lectin domains. Representation of the VP8* domain of SA11 (PDB 1KQR) coloured according to its secondary structure. The glycan binding sites located in the cleft between the β-sheets (I) and adjacent to it (II) are indicated. (B-D) Accessibility of sites I (green line) and II (blue line) in the head (B, C) and stem (D) lectin domains of the NTR (B, D) and TR (C) spike.

**Figure S8.**
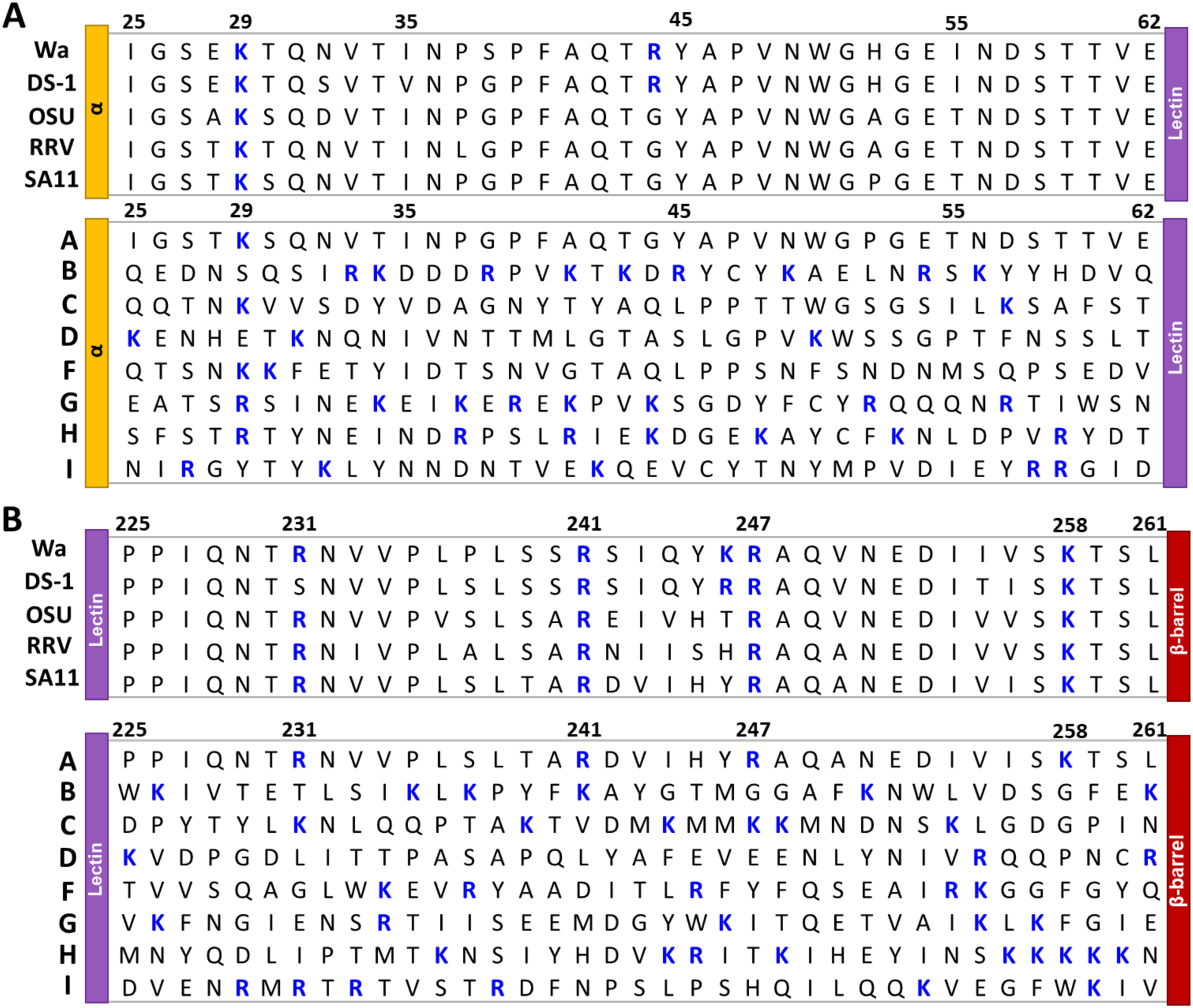
Multiple alignment of the amino acid sequences of protein VP4 from different RV strains in the regions surrounding the trypsin digestion sites. **(A)** Sequences corresponding to the α2-β1 loop (between the α and lectin domains, residues 25 to 35). **(B)** Sequences corresponding to the α3− β14 loop (between the lectin and β-barrel domains of the spike body, residues 225 to 261). The sequences shown correspond to representative strains of the RVA species (top panel) and to the reference strain of the different RV species (bottom panel). Residues susceptible to being cut by trypsin, lysine (K) and arginine (R), are marked in blue. All aa are presented with the single letter code.

## Supplementary table

**Table S1.**
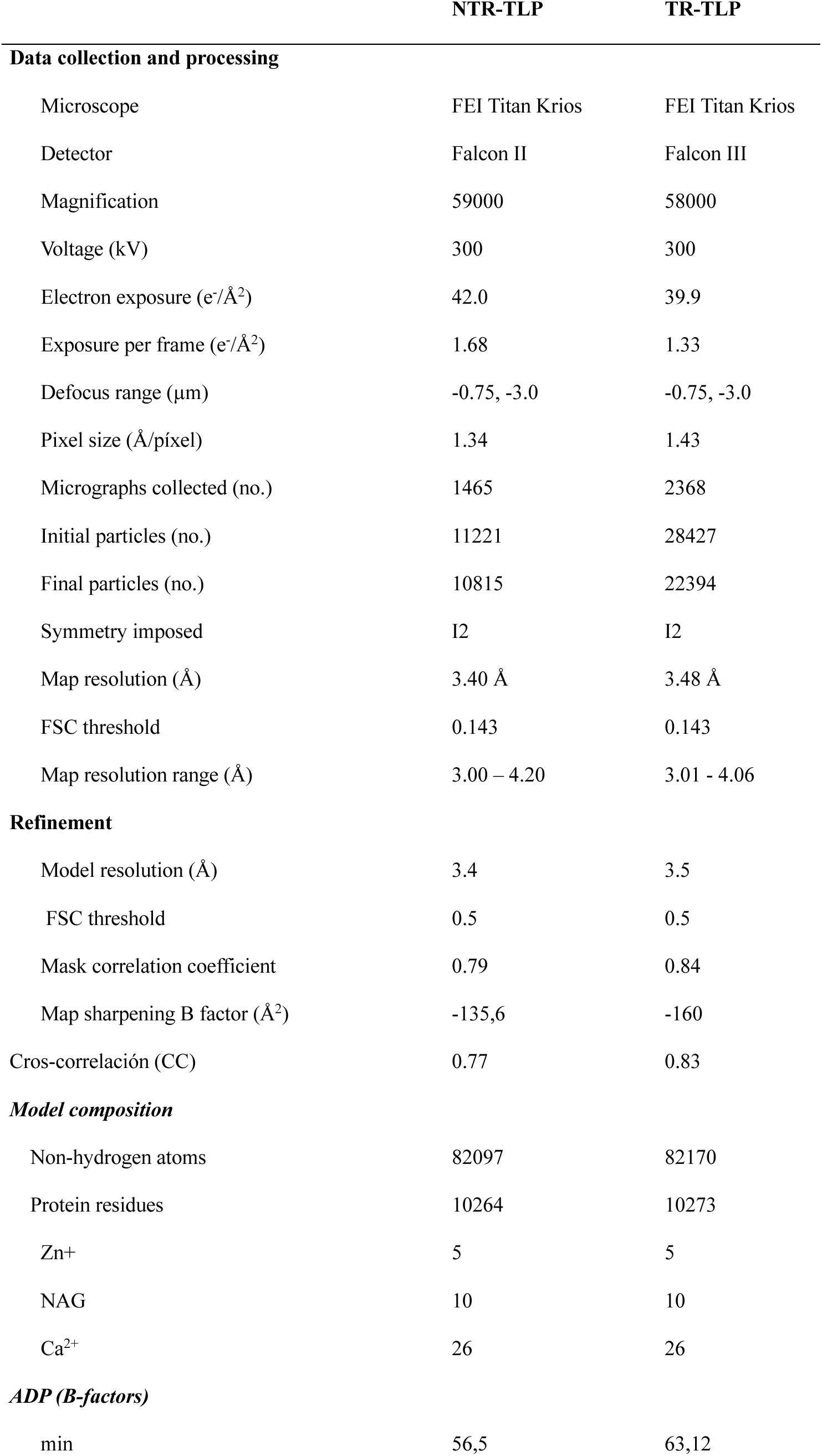
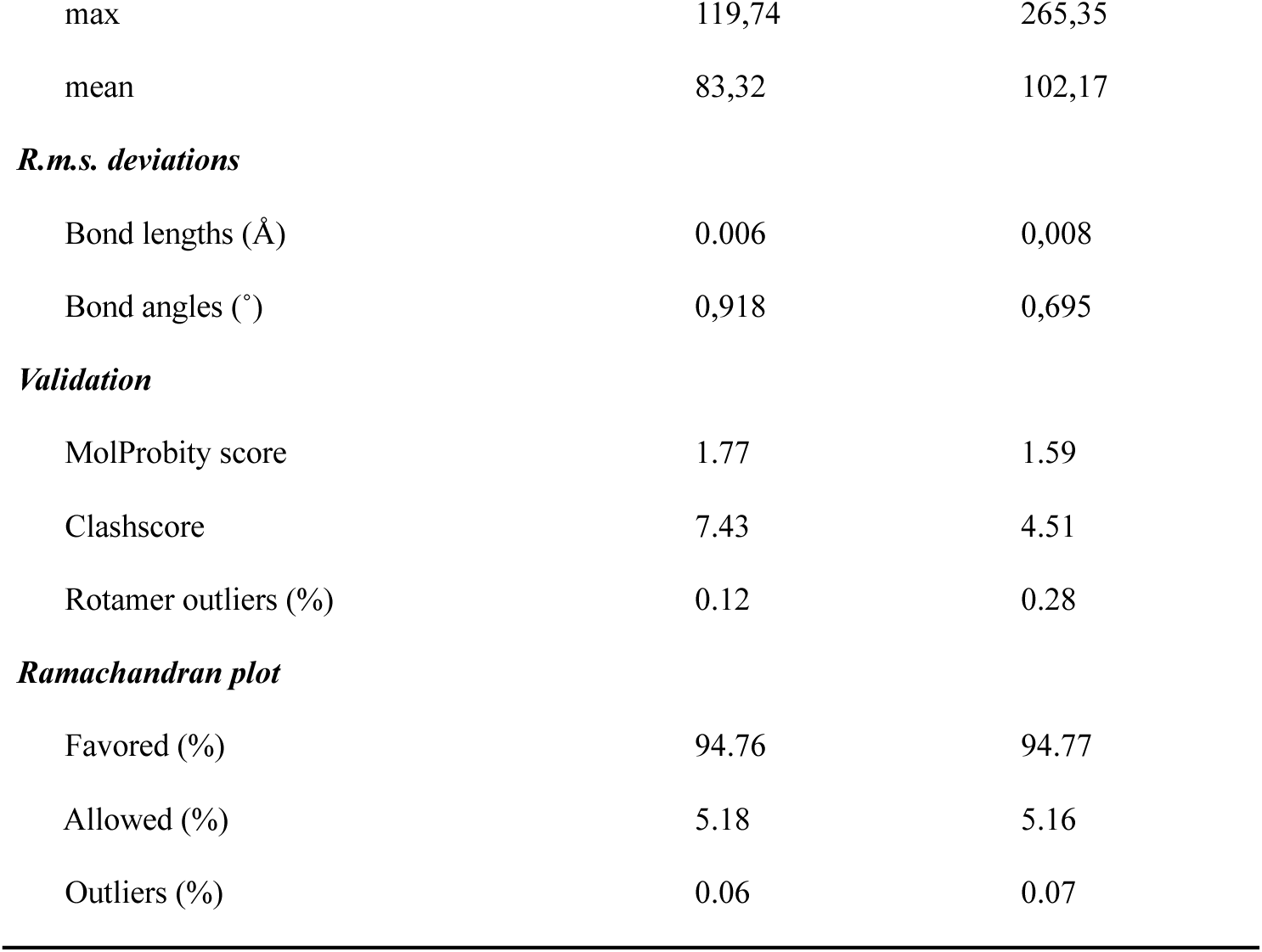
Cryo-EM data collection and model statistics.

